# Bacteriophages Phi 8 and Phi 12 host infection are inhibited by OMVs and LPS purified from *P. pseudoalcaligenes* strain: East River Isolate A

**DOI:** 10.1101/2024.09.02.610742

**Authors:** Charles F. Robinson, Reza Khayat

## Abstract

*Cystoviridae* is a family of double stranded RNA (dsRNA) phage that infects various strains of *Pseudomonas syringae*, a Gram-negative soil bacteria known to infect various crops. Surrounding the icosahedral capsids of these phages is a bacterial derived phospholipid membrane. Embedded within this membrane is a multi-component protein complex, referred to as the spike complex. The spike complex is responsible for host recognition and membrane fusion. We studied the ability of two members of the *Cystivirdae* family to infect cells in the presence of purified outer membrane vesicles (OMVs) and lipopolysaccharide (LPS) derived from distinct sources. In this study we determined that OMVs from the host *Pseudomonas pseudoalcaligenes* strain: East River isolate A (ERA) inhibit Phi 8 and Phi 12 host infection. These OMVs range in size from 30 to 60 nm and bind to Phi 8 and Phi 12. However, OMV purified from *P. syringae* pv. phaseolicola LM2691 and *E. coli* Δ*yciB* Δ*dcrB* did not inhibit Phi 8 or Phi 12 host infection. However, LPS derived from ERA and LM2691 inhibited Phi 8 and Phi 12 infection, demonstrating that LPS is the receptor for these two viruses, and that OMV biogenesis is selective of LPS. LPS derived from other non*-Cystoviridae* Gram-negative bacteria, did not inhibit infection. We confirmed that host proteins are not required for Phi 8 or Phi 12 host interaction. Our results also suggest that differences in lipid A and the core polysaccharide in LPS may influence Phi 8 and Phi 12 host binding.

**IMPORTANCE:** Most phage families studied to date use a tailed appendage, composed of a multitude of proteins, for cellular recognition, membrane penetration, and genome injection. This contrasts with members of the *Cystoviridae* family which possess a phospholipid membrane bilayer with embedded proteins responsible for cellular recognition and membrane fusion. Thus, the *Cystoviridae* are akin to enveloped viruses which also use protein complexes embedded into their membrane for cellular recognition and membrane fusion. Examples of such viruses include the *Retroviridae, Coronoviridae, Herpesviridae*, and *Orthomyxoviridae* families. The binding specifics of *Cystoviridae* to the host outer membrane are unknown. Using *Cystoviridae*-OMV interaction we began to uncover the host requirements for binding *Cystoviridae*. The results presented determine that only lipid A and the core polysaccharide of LPS are required for *Cystoviridae* outer membrane binding.

## INTRODUCTION

The *Cystovirdae* family is a bacteriophage with an external phospholipid bilayer membrane surrounding an icosahedral nucleocapsid [2]. Within the capsid is the genome composed of three segments of dsRNA. These segments are named the large segment (L, 6.4 kbp), the medium segment (M, 4.1 kbp), and the small segment (S, 2.9 kbp) [3]. Embedded into the phospholipid envelope is protein P6, and attached to this protein is P3. Together P3 and P6 are responsible for binding and fusing with the outer membrane of the plant pathogen host *Pseudomonas syringae* [4]. Following membrane fusion, the P5 protein hydrolyzes the peptidoglycan of the host so that the phage can gain entry into the host’s cytoplasm [5]. Within the cytoplasm, the *Cystovirdae* RNA is then transcribed by P2 into sense mRNA, which is then translated by the host ribosomes. The polymerase complex, P1, is assembled after translation. P1 also acts as the major capsid protein and forms the procapsid. The RNA is then reinserted into the procapsid by P4 and replicated by P2. The icosahedral lattice nucleocapsid protein P8 forms around the procapsid. P9 generates the envelope around the nucleocapsid with the help of the proposed chaperone P12. Once the complete virion is assembled it lysis the host cell [6].

Two members of the *Cystovirdae* family are Phi 8 and Phi 12. They were first identified by isolation from leaves of plants and genetically sequenced [7, 8]. Phi 8 has a heterodimer P3 composed of P3a and P3b. While Phi 12 has a heterotrimer composed of P3a, P3b, and P3c [9, 10]. Cryo-electron tomography studies of Phi 12 have visualized the structure of P3. It was determined that P3 adopts a hexameric oligomer that is approximately 19 nm in length [11]. P3 is attached to P6 to form the spike complex [4]. It is known that Phi 8 and Phi 12 use P3 to bind directly to the rough LPS of *P. syringae* [12]. We investigated what the host membrane requirements to bind Phi 8 and Phi 12.

Gram-negative bacteria possess an asymmetric outer membrane that contains LPS in the outer leaflet. At the extremity of LPS, furthest from the membrane, are O-antigen polysaccharides, which are then attached to the core polysaccharides, and then lipid A. The O-antigen consists of repeating units of oligosaccharides. The core polysaccharide contains sugars 3-deoxy-D-mannooctulosonic acid (KDO) and hexoses. Lipid A is an amphipathic lipid containing glucosamine, acyl chains and phosphate groups [13]. Neighboring LPS molecules share numerous hydrogen bonds between their saccharides, divalent cations bridged between their KDO, and hydrophobic packing between their acyl chains to generate strong lateral interactions. These interactions are responsible for preventing the transportation of small hydrophobic and hydrophilic molecules across the LPS [14]. Most phages (e.g. those from the *Siphoviridae, Myoviridae, Podoviridae*, and *Microviridae* family) overcome this barrier by using a syringe-like appendage that injects their genomes through the LPS of their hosts. Only the genome enters the host while the other phage machinery remains outside [15]. The members of the *Cystoviridae* family are unique in that they fuse their phospholipid outer membrane with the LPS of the host and their entire capsid enters the host [1].

Gram-negative bacteria, including *P. syringae*, can naturally secrete their outer membrane in the form of OMVs [16]. These vesicles contain LPS, proteins, and carbohydrates moieties on their outer leaflet. OMVs are believed to be generated to relieve environmental stress [17]. Also, OMVs act as a decoy for phage infection. The OMVs neutralize the phage by binding to the host binding protein of the phage and prevent infection [18]. In this study, we used plaques assays with purified OMVs and LPS to determine the host requirements for binding Phi 8 and Phi 12. In particular, we purified OMVs and LPS from *P. syringae* pv. phaseolicola LM2691 and *Pseudomonas pseudoalcaligenes* strain: East River isolate A. Utilizing plaque assays we determined that purified OMVs from *Pseudomonas pseudoalcaligenes* strain: East River isolate A inhibited both Phi 8 and Phi 12 host infection. While purified LPS from *P. syringae* pv. phaseolicola LM2691 and *Pseudomonas pseudoalcaligenes* strain: East River isolate A inhibited Phi 8 and Phi 12 host infection. The interaction between *Cystovirdae*-OMV/LPS was then visualized with negative staining electron microscopy. We demonstrate that Phi 8 and Phi 12 are more efficient at binding to purified LPS from ERA. This study begins to uncover the identity of Phi 8 and Phi 12 host receptors.

## MATERIALS AND METHODS

### Bacterial Strains

Phi 8 and Phi 12 host strains used were *P. syringae* pv. phaseolicola LM2691 (LM2691) [19] and *Pseudomonas pseudoalcaligenes* strain: East River isolate A (ERA) [20]. LM2691 is a rough LPS derivative strain of LM128. The strain LM128 is a derivative strain of the smooth LPS strain HB10Y. LM2691 contains the plasmid pLM10866. This plasmid expresses a T7 RNA polymerase [19]. ERA is a rough LPS alternative host for the *Cystovirdae* family [21]. The OMV producing mutant *E. coli* Δ*yciB* Δ*dcrB* (Strain A1139) was utilized as a negative control [22]. Host cells were stored at −80°C in 80% (v/v) glycerol and plated on Luria-Bertani (LB) agar at 25°C for 36 hours. Host were grown in LB broth at 25°C. The host was then infected at an OD_600_ of 1.2.

### Phage Preparation

A 1 L host culture was grown to stationary phase (OD_600_=1.2) and centrifuged at 5,000 x g for 15 minutes at 4°C. The pellet was then resuspended in 50 ml of LB broth. The resuspended host was then incubated with Phi 8 or Phi 12 for 15 minutes. The mixture was then incubated with top agar (0.8% (w/v) agar) and plated on LB agar plates. Plates were incubated for 16 hours at 25°C. The top agar was then scraped off the LB agar and centrifuged for 26,000 x g at 4°C for 1 hour (Beckman Coulter JA-17). The supernatant was then removed, and the centrifuge step was repeated. The supernatant was then spun at 100,000 x g for 1 hour (Beckman Coulter Type 45 Ti) and resuspended in P buffer (10 mM potassium phosphate, 1mM MgCl_2_, pH 8.0). The resuspended pellet was then placed on a sucrose gradient of 10% (w/v) - 30% (w/v) for 1 hour at 63,400 x g at 4°C. The 1 ml fraction containing the phage was then placed on a 40% (w/v) - 60% (w/v) sucrose gradient for 1 hour at 63,400 x g at 4°C (Beckman Coulter SW 40 Ti). The 1 ml fraction was then placed in a 50 kDa dialysis membrane (Spectrum Labs) in a buffer of 20 mM KH_2_PO_4_, 50 mM NaCl, pH 7.5 for 16 hours. The phage was then removed and placed in a 1 kDa dialysis membrane (Spectrum Labs) in a buffer of 10% (w/v) PEG 3350, 20 mM KH_2_PO_4_, 50 mM NaCl, pH 7.5 until concentrated to 500 μl. The Phage was stored in −80°C in 80% (v/v) glycerol. Phi 8 and Phi 12 had a PFU of 3.4×10^11^ PFU/ml and 7.8×10^10^ PFU/ml, respectively [23]. PFU values when pertaining to the host *Pseudomonas pseudoalcaligenes* ERA differed for Phi 8 and 12 at values of 8.2×10^10^ PFU/ml and 5.8×10^5^ PFU/ml, respectively.

### OMV Purification

Cultures of 1 L host were grown in LB broth at 25°C for 72 hours (OD_600_ = 2.0). The host was then pelleted at 5,000 x g for 10 minutes at 4°C. The supernatant was then filtered and concentrated using a Minimate Tangential Flow Filtration system with a molecular weight cutoff of 500 kDa (Pall Scientific) [24]. The sample was concentrated to a volume of 10 ml and then buffer exchanged with 100 ml of phosphate buffered saline (137 mM NaCl, 2.7 mM KCl, 10 mM Na_2_HPO_4_, 1.8 mM KH_2_PO_4_, pH 7.4) [25]. The sample was further concentrated using a spinning filter with a 100 kDa molecular weight cutoff (Pall Scientific) spun at 4,500 x RPM (Beckman Coulter SX4250) for 40 minutes at 4°C.

### LPS Isolation

*Escherichia coli* (*E. coli*) F583 (Lot 128M4131V), *Pseudomonas aeruginosa* (*P. aereginosa*) 10 (0000105248), and *Salmonella enterica* (*S*. enterica) Minnesota Re 95 rough (0000138313) LPS were ordered from Millipore Sigma. LPS was resuspended in distilled water to a concentration of 3 mg/ml. LM2691 and ERA LPS were isolated using the Darveau-Hancock Method [26]. A 1 L culture of cells were grown to an OD_600_ of 1.0 in LB broth and then pelleted at 7,500 x RPM (Beckman Coulter JLA-8.1000) for 15 minutes at 4°C. The pellet was resuspended with 15 ml of 10 mM Tris-HCl (pH 8.0) and 2 mM MgCl_2_. Extracellular nucleic acids were digested with DNase (100 μg/ml) and RNase (25 μg/ml).The sample was then lysed with two cycles of the French press at 15,000 psi. The sample was further lysed with macro tip sonication by two pulses at an intensity of 30% (34 W) for 30 seconds. Further nucleic acid digestion was performed with DNase and RNase at final concentrations of 200 μg/ml and 50 μg/ml, respectively. The sample was then incubated on a shaker at 37°C for 2 hours. Once cooled to room temperature 5 ml of 0.5 M EDTA (tetra sodium salt), 10 mM Tris-HCl (pH 8.0); 2.5 ml of 20 % (w/v) SDS, 10 mM Tris HCl (pH 8.0); and 2.5 ml of 10 mM Tris-HCl (pH 8.0) were added and adjusted to a pH of 9.5. The sample was then vortexed and ultracentrifuged at 50,000 x g for 30 minutes at 20°C (Beckman Coulter Type 45 Ti). Proteinase K (200 μg/ml) was added to the supernatant and incubated on a shaker at 37°C overnight. Two volumes of 0.375 M MgCl_2_ in 95% ethanol were combined with the sample and cooled to 0°C. The cooled mixture was then centrifuged at 12,000 x g for 15 minutes at 4°C. The pellet was then resuspended in 25 ml of 0.1 M EDTA, 2% (w/v) SDS, 10 mM Tris-HCl (pH 8.0). The resuspended pellet was then lysed with a sonicator macro probe at an intensity of 30% for two bursts (30 seconds). The solution was then incubated at 85°C for 30 minutes and cooled to room temperature. The pH of the solution was then adjusted to 9.5 with 4M of NaOH. Proteinase K was then added to a final concentration of 25 μg/ml. The solution was incubated at 37°C overnight with constant shaking. Two volumes of 0.375 M MgCl_2_ in 95% ethanol were added to the solution and cooled to 0°C. The mixture was then centrifuged at 12,000 x g for 15 minutes at 4°C. The pellet was resuspended in 15 ml of 10 mM Tris-HCl (pH 8.0) and sonicated at an intensity of 30 for two bursts at 30 seconds. The solution was then centrifuged at 1,000 x RPM (Beckman Coulter SX4250) for 5 minutes. The supernatant was obtained, and the pellet was washed with 500 μl of distilled water. The resuspended pellet was then spun at 1,000 x RPM (Beckman Coulter SX4250) for 5 minutes. The supernatant of the washed pellet was then combined with the original supernatant from the previous centrifugation. The solution was then given a final concentration of 25 mM MgCl_2_ and ultracentrifuged at 200,000 x g for two hours (4°C) (Beckman Coulter Type 45 Ti). The pellet was then resuspended in 100 μl of distilled water.

### KDO Quantification

The concentration of KDO was quantified to determine the concentration of purified LPS and concentration of LPS in OMV. The KDO concentration was quantified according to Sunayana and Reddy 2015 [27]. A 50 μl sample of purified OMV or LPS was added to 50 μl of 0.5 N H_2_SO_4_. The mixture was then heated to 100°C for 8 minutes. Once cooled 50 μl of periodate (2.28×10^-3^ g/ml) was then added and incubated for 10 minutes at room temperature. After incubation, 200 μl of sodium arsenite (4.0×10^-2^ g/ml in 0.5 N HCl) and 800 μl of thiobarbituric acid (6 g/ml) was then added to the mixture and heated at 100°C for 10 minutes. After cooling, 1.5 ml of 95% (v/v) n-butanol in concentrated HCl was added and then spun at 2,000 x RPM (Beckman Coulter SX4250) for 5 minutes at room temperature. The upper butanol layer was then aspirated. The difference in optical density at 552 nm and 509 nm was obtained to determine the concentration of KDO (19 = 1 μM).

### Plaque Assays

The host was grown overnight in LB broth at 25°C. A fresh subculture was then standardized to an OD_600_ of 0.1 using the overnight culture. The culture was then grown to an OD_600_ of 1.2. Phi 8 or Phi 12 was then added to 600 μl of host for 15 minutes. Top agar (0.8 % (w/v) agar) was then added to the host-phage mixture. The mixture was then plated on LB agar at room temperature. Plaques were quantified after 16 hours. For plaque assays with OMVs or LPS, phage was incubated with OMVs or LPS at a KDO concentration of 3 μg/ml, 300 ng/ml, or 3 ng/ml at 4°C on a revolver for 40 minutes. To normalize the number of plaques, Plaque forming units (PFU) per ml and Log (treated PFU/initial PFU) were determined.

### Transmission Electron Microscopy of Phage-OMV/LPS interaction

Phage and LPS or OMV (1:4) were incubated for 40 minutes at room temperature. The mixture was then placed on a glow discharged carbon coated 400 mesh copper grid for 30 seconds. The sample was then stained with 2 % (w/v) uranyl formate for 10 seconds and then 30 seconds. Images were obtained with a JEOL 2100 TEM at 200 kV. Images were captured at 40,000x magnification and 12,000x magnification.

### Statistical Analyses

Graphs and statistical analysis were generated using GraphPad Prism software. P values were determined using one-way analysis of variance (ANOVA).

## Math and equations

### Plaque Forming Units (PFU)

Plaques were normalized by determining the plaque forming units per ml.

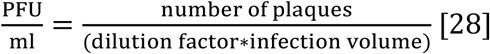

### Logarithmic Change in PFU Treated with Purified OMVs or LPS

PFU treated with OMVs or LPS log change were determined by the equation below:

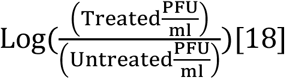

## RESULTS

### OMV derived from *P. pseudoalcaligenes* ERA inhibit Phi 8 and Phi 12 infection

We asked, can OMV derived from hosts of Phi 8 and Phi 12 mimic the host to the extent that they inhibit infection? To answer this question, we performed plaque assays in the presence of increasing concentration of OMV purified from ERA and LM2691. Both Phi 8 and Phi 12 infect the LM2691 and ERA cell lines. Increasing concentrations of OMV derived from ERA cells inhibited plaque formation by both Phi 8 and Phi 12. To estimate the extent of inhibition, we measured the logarithmic change in plaque formation (PFU) due to the OMV (**Fig. 1)**. The results clearly demonstrate that ERA derived OMV diminish infection of Phi8 and Phi 12 by two orders of magnitude. Thus, OMV derived from ERA cells are indistinguishable from the host to both Phi 8 and Phi 12. However, OMV derived from LM2691 slightly inhibited Phi 8 and Phi 12 infection of LM2691 host cells. This indicates that OMV derived from LM2691 are sufficiently distinct from the cells for Phi 8 and Phi 12 to differentiate.

**FIG 1.**
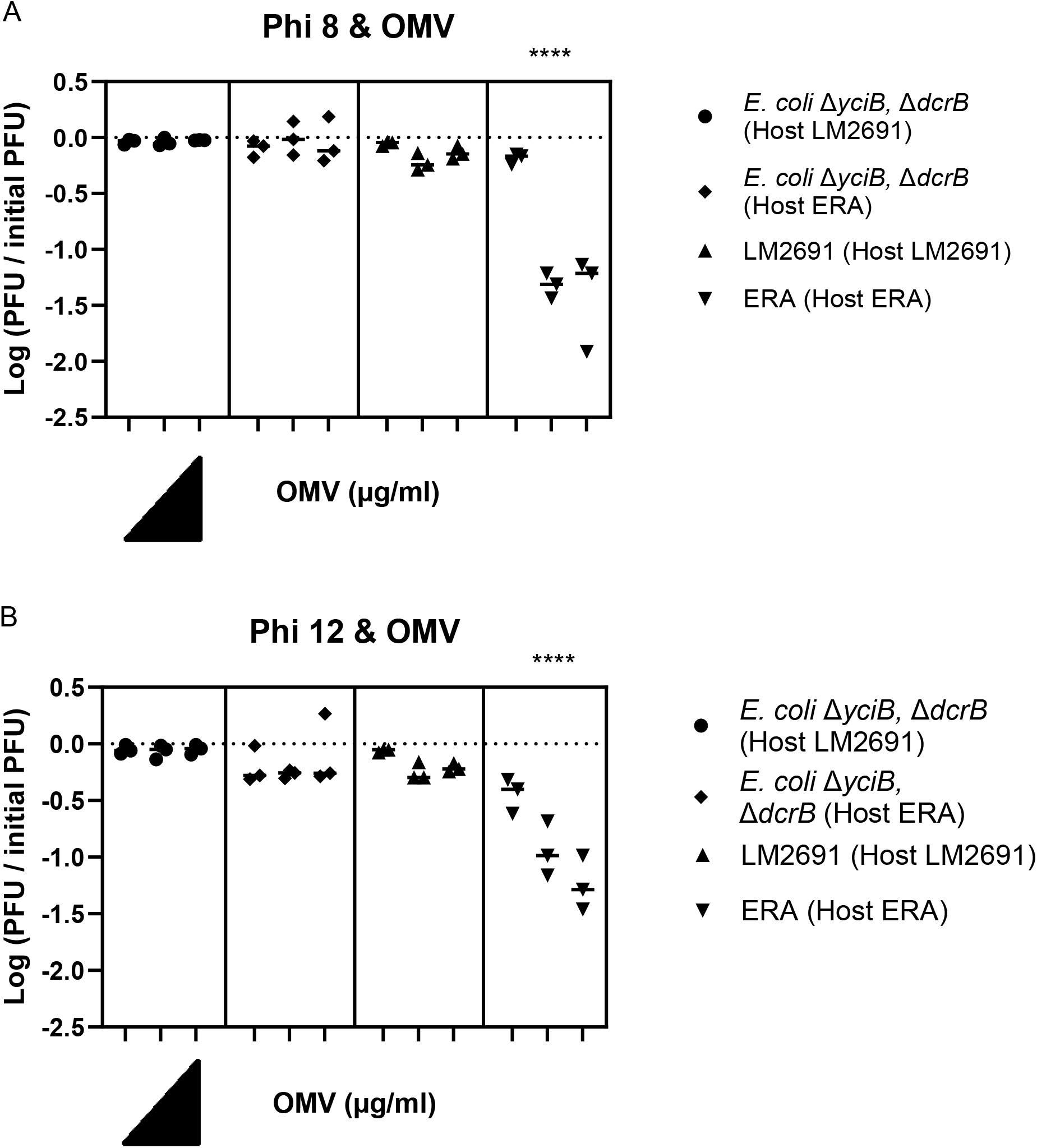
Inhibition of Phi 8 and Phi 12 by OMV. OMV derived from *Pseudomonas pseudoalcaligenes* East River isolate A (ERA) inhibits Phi 8 (of 2.74 x 10^11^ PFU/ml) and Phi 12 (8.1 x 10^10^ PFU/ml) infection of host LM2691 or ERA (Paratheses). Host was grown to 2.5 x 10^9^ CFU/ml and infected with A. Phi 8 or B. Phi 12. KDO concentrations of 3 ng/ml, 300 ng/ml, or 3 μg/ml (Left to right) were added to host-phage mixture; n = 3. Solid lines indicate mean. ^***^, P ≤ 0.001; ^****^, P ≤ 0.0001

We then asked, can OMV purified from a non-infectious Gram-negative bacteria inhibit Phi8 and Phi12 from infecting ERA or LM2691 cells? To answer this question, we purified OMV from a hyper vesiculating *E. coli* strain Δ*yciB* Δ*dcrB*. This strain has an altered membrane function that leads to an immature toxicity of the outer membrane lippoprotein Lpp [22]. As anticipated, OMV from *E. coli* Δ*yciB* Δ*dcrB* did not inhibit infection of ERA or LM2691. This data demonstrates that OMV purified from a non-permissive Gram-negative strains are unable to inhibit Phi 8 and Phi 12 infection (**Fig. 1**).

### LPS derived from *P. syringae* LM2691 and *P. pseudoalcaligenes* ERA inhibit Phi 8 and Phi 12 infection

We then asked if LPS purified from hosts, and non-hosts were sufficient to inhibit Phi 8 and Phi 12 infection of LM2691. To answer this question, we performed plaque assays in the presence of increasing concentrations of LPS. LPS purified from LM2691, *E. coli, P. aereginosa, S. enterica*, and ERA were incubated with Phi 8 or Phi 12 for 40 minutes. The phage-LPS mixture was then incubated with the host for 15 minutes. Plaque assays were then performed. The results indicate that LPS purified from *P. syringae* LM2691 and *P. pseudoalcaligenes* ERA are sufficient to inhibit Phi 8 and Phi 12 infection. While *P. aereginosa* LPS did exhibit a consistent yet slight inhibition of infection, statistical analysis indicates this to be insignificant. Non Cystoviridae host LPS were not able to inhibit Phi 8 or Phi 12 infections (**Fig. 2**).

**FIG 2.**
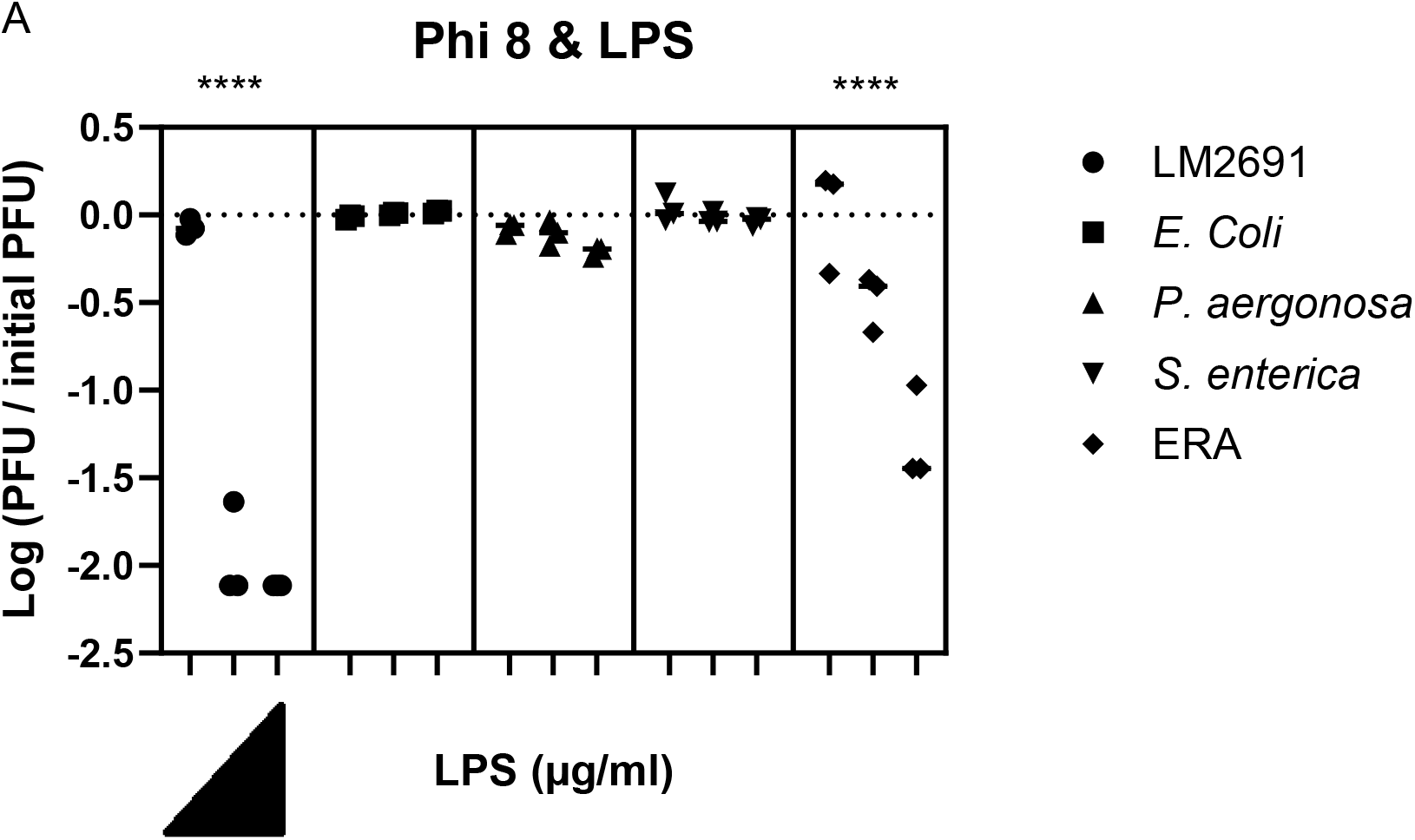
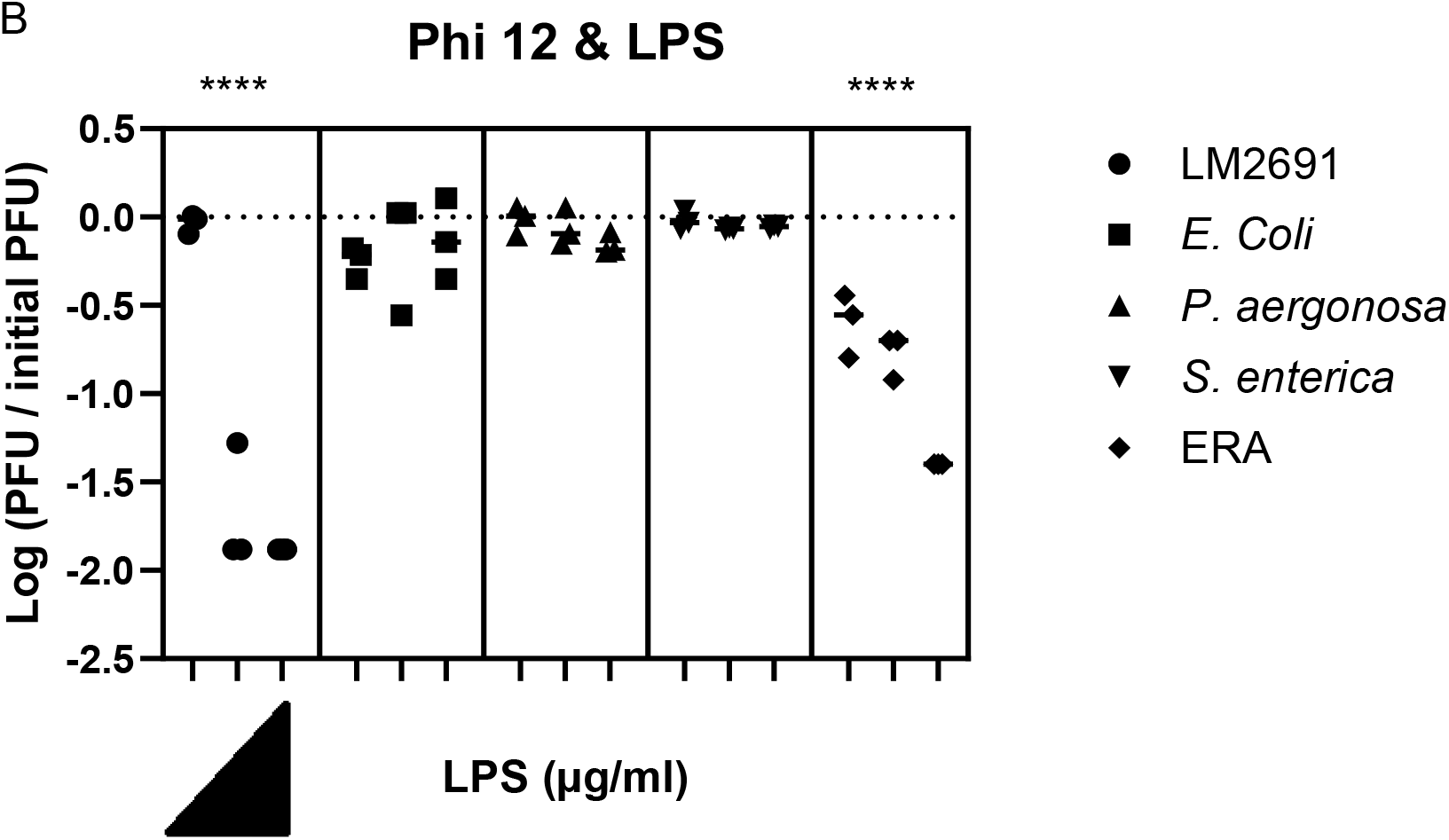
Inhibition of Phi 8 and Phi 12 by purified LPS. Purified LPS from *Pseudomonas syringae* LM2691 inhibits A. Phi 8 (of 1.3 x 10^10^ PFU/ml) and B. Phi 12 (7.6 x 10^9^ PFU/ml) infection of host *Pseudomonas syringae* LM2691. Host LM2691 was grown to 2.5 x 10^9^ CFU/ml and infected. KDO concentrations of 3 ng/ml, 300 ng/ml, or 3 μg/ml (Left to right) were added to host-phage mixture; n = 3. Solid lines indicate mean. ^***^, P ≤ 0.001; ^****^, P ≤ 0.0001

### Phi 8 and Phi 12 bind to the *P. pseudoalcaligenes* ERA OMV & LPS and *P. syringae* LM2691 LPS

Having demonstrated that OMV purified from *P. pseudoalcaligenes* ERA inhibit Phi 8 and Phi 12 infections, we asked if we could visualize the interaction between the OMV and phage. We visualized the interaction between phage and OMV using negative stained electron microscopy (NSEM). *P. pseudoalcaligenes* ERA OMVs were incubated with Phi 8 or Phi 12 and then placed on a carbon coated grid, stained with uranyl formate, and imaged with a transmission electron microscope (TEM). OMV sizes ranged from 10 nm to 80 nm. Phi 8 and Phi 12 have an average size of ∼92 nm. The *P. pseudoalcaligenes* ERA OMV bound to Phi 8 and Phi 12 were of sizes 30 nm to 60 nm (n=4). Only one OMV was found to be bound to one Phi 8 particle (n=3) and two bound to one Phi 12 particle (n=4). Thus, our results suggest that Phi 8 and Phi 12 binds to *P. pseudoalcaligenes* ERA OMV to inhibit host infection (**Fig. 3**).

**FIG 3.**
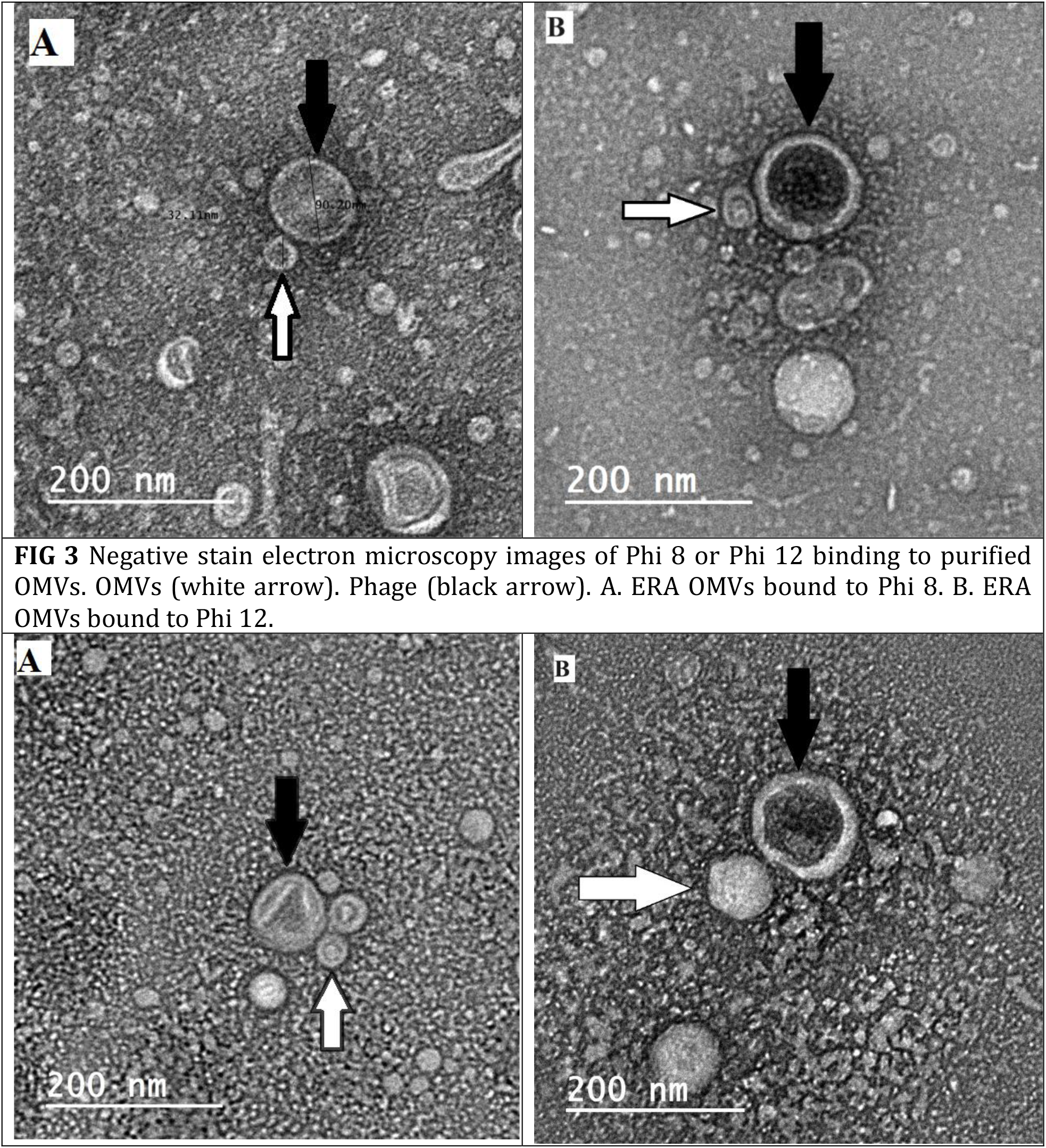
Negative stain electron microscopy images of Phi 8 or Phi 12 binding to purified OMVs. OMVs (white arrow). Phage (black arrow). A. ERA OMVs bound to Phi 8. B. ERA OMVs bound to Phi 12.

We then wanted to determine how purified LPS interacts with Phi 8 and Phi 12. NSEM was performed on Phi 8 and Phi 12 preincubated with purified LPS from ERA and LM2691. TEM images indicate that LPS purified from both host strains form vesicles that range from 30 nm to 90 nm in size (n=15). LPS morphology also varies from spherical to non-spherical. LPS derived from *P. syringae* LM2691 or *P. pseudoalcaligenes* ERA was found to be bound to Phi 8 and Phi 12. The phage bound LPS ranged from 20 nm to 50 nm (n=3). As many as three LPS vesicles were found to bind to the phage. This also suggests that *P. syringae* LM2691 and *P. pseudoalcaligenes* ERA LPS binds to Phi 8 and Phi 12 to inhibit infection (**FIG. 4**.).

**FIG 4.**
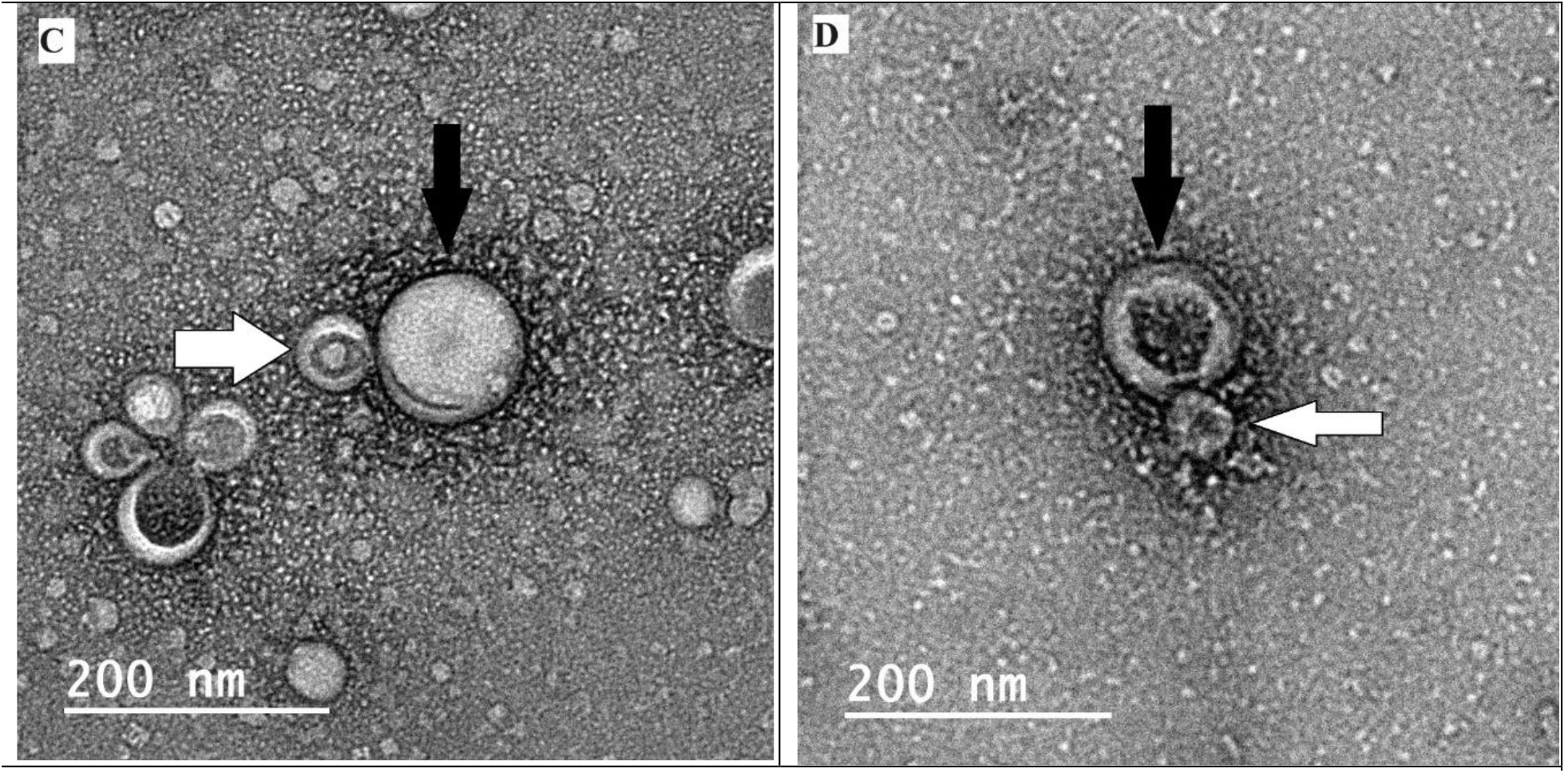
Negative stain electron microscopy images of Phi 8 or Phi 12 binding to purified LPS. LPS (white arrow). Phage (black arrow). A. ERA LPS bound to Phi 8. B. LM2691 LPS bound to Phi 8. C. ERA LPS bound to Phi 12. D. LM2691 LPS bound to Phi 12.

## DISCUSSION

The RNA phage family *Cystoviridae* has the unique ability to fully enter the bacteria host. This is accomplished by P3 binding to the rough LPS of the host. Then, P6 fuses with the outer membrane of the host [4]. Currently the specifics of *Cystoviridae* binding are unknown.

Plaque assays performed in the presence of purified components can provide insight into viral entry. OMVs are host derived and can contain LPS and outer membrane proteins, while purified LPS lacks proteins [29]. Previously, researchers have determined that rough LPS is required for Phi 8 and Phi 12 host binding. This was done by comparing plaque formation between rough and smooth LPS host. [12]. Also, researchers have determined that the *Salmonella typhimurium* phage P22 and the *Vibrio Cholera* phage ICP infections are inhibited by host OMVs. These phages that are inhibited by OMVs contain tails for attachment. [18, 30]. Unlike *Cystoviridae* that contains a spike complex [4]. We set to demonstrate the binding specifics of Phi 8 and Phi 12 to the host membrane by studying OMV-Phage interaction. In this study we demonstrate that LPS alone is sufficient to bind Phi 8 and Phi 12.

Our results demonstrate that the ability of Phi 8 and Phi 12 to infect their host is significantly inhibited in the presence of purified OMVs and LPS from *P. syringae* LM2691 and *P. pseudoalcaligenes* ERA. Phi 8 and Phi 12 inhibition were comparable to each other. We confirm that different *Cystoviridae* host can not only produce OMVs but significantly inhibit phage infection.

Researchers have also found that the presence of the O antigen may influence the shape of purified LPS. Purified Smooth type LPS from *E. coli* was found to have filamentous shape. While purified rough LPS from *E. coli* was less filamentous [31]. We determined the size of purified OMVs ranged from 30 nm to 80 nm. Shapes varied but the OMVs that bound to Phi 8 and Phi 12 were mostly spherical. Purified LPS had a smaller range from 20 nm to 50 nm. A majority of purified LPS was spherical with no filamentous shape being observed. Our negative staining TEM images suggest that Phi 8 and Phi 12 binding to OMV can be correlated to inhibition of infection.

The *Cystoviridae* host ranges from *P. syringae, P. savastanoi*, and *P. pseudodoalcaligenes* [9, 32]. Plaque assays with the host *P. pseudodoalcaligenes* ERA had a decrease in PFU when compared to *P. syringae* LM2691. Plaques also had a decrease in size when compared to plaques formed by LM2691. These differences in infection of hosts may contribute to the difference of OMV and LPS inhibition of infection. OMV derived from LM2691 did not inhibit phage infection, but its LPS form did. This occurrence may be due to difference in purification methods, OMVs lacking rough LPS, or obstruction of the LPS receptor by being embedded into the OMV. LPS purification is obtained by lysing the host. While OMV purification only isolates sections of the host. Thus, the OMVs produced by the host may lack the *Cystovirdae* receptor. *E. coli* JM109 can be infected by Phi 8 and Phi 12 but will not form plaques. *E. coli* JM109 has partial truncated rough LPS that will allow Phi 8 and Phi 12 to kill it, but not form plaques [12]. *E. coli* Δ*yciB* Δ*dcrB* strain is a derivative of MG1655 not JM109 [22]. *E. coli* Δ*yciB* Δ*dcrB* is known to over vesiculate [22]. We attempted OMV purification of JM109 but no OMVs were isolated. In conclusion, the purified rough LPS of *E. coli* Δ*yciB* Δ*dcrB* could not inhibit Phi 8 or Phi 12 infections.

Previously, researchers have investigated the structure of the *P. syringae pv*. phaseolicola rough LPS. The core polysaccharide is defined by N-acetylglucosamine and rhamnose. The outer core polysaccharide alternates between two glycoforms [33]. The lipid A of *P. syringae pv*. phaseolicola contains glucosamine with two or four phosphate groups bound to it. Also, glucosamine has primary and secondary acyl chains bound to it with hydroxylation occurring on both chains. *P. aeruginosa* has a similar inner core polysaccharide with varying degrees of phosphorylation. The outer core of *P. aeruginosa* differs from *P. syringae* by the terminal rhamnose position. Lipid A of *P. aeruginosa* also contains glucosamine but differs in the number of phosphate groups and hydroxylation [34]. The differences and similarities in LPS structure between *P. syringae pv*. phaseolicola and *P. aeruginosa* may contribute to the weak binding of the *Cystovirdae* to purified *P. aeruginosa* LPS.

## ACKNOWLEDGEMETS

We thank the Anuradha Janakiraman lab for the *E. coli* strain *E. coli* Δ*yciB* Δ*dcrB* (Strain A1139). We thank CCNY Microscopy Facility for access to the instrumentation needed for these studies.

